# Inorganic and black carbon hotspots constrain blue carbon mitigation services across tropical seagrass and temperate tidal marshes

**DOI:** 10.1101/2020.10.02.310946

**Authors:** John Barry Gallagher, Vishnu Prahalad, John Aalders

## Abstract

Total organic carbon (TOC) sediment stocks as a CO_2_ mitigation service requires exclusion of allochthonous black (BC) and particulate inorganic carbon corrected for water– atmospheric equilibrium (PIC_eq_). For the first time, we address this bias for a temperate salt marsh and a coastal tropical seagrass in BC hotspots. Seagrass TOC stocks were similar to the salt marshes with soil depths < 1 m (59.3 ± 11.3 and 74.9 ± 18.9 MgC ha^-1^, CI 95% respectively) and sequestration rates of 1.134 MgC ha^-1^ yr^-1^. Both ecosystems showed larger BC constraints than their pristine counterparts. However, the seagrass meadows’ mitigation services were largely constrained by both higher BC/TOC and PIC_eq_/TOC fractions (38.0% ± 6.6% and 43.4% ± 5.9%, CI 95%) and salt marshes around a third (22% ± 10.2% and 6.0% ± 3.1% CI 95%). The results demonstrate a need to account for both BC and PIC within blue carbon mitigation assessments.

## 1. Introduction

Blue carbon refers to the carbon sequestered and stored in salt marsh, mangrove, and seagrass beds (Nellemann et al., 2009). There has been a growing interest in measuring, mapping, and valuing blue carbon stocks to promote their importance in climate change mitigation (Kelleway et al., 2017; Serrano et al., 2019). However, the use of total organic carbon (TOC) for sequestration and stock assessments has recently been in question. Traditional assessments have failed to remove allochthonous organic recalcitrants from the equation (Chew and Gallagher, 2018; Chuan et al., 2020; Gallagher, 2014; Gallagher, 2015; Gallagher, 2017; Gallagher et al., 2020 in press; Gallagher et al., 2019). These stable organic forms are not produced by the blue carbon ecosystem and deposition by the ecosystem does not afford additional protection from remineralization. Consequently, their presence is not a measurable blue carbon service of storage or sequestration in the mitigation of greenhouse gas emissions. The argument is unequivocal and has now been established as an important blue carbon constraint by the IPCC (Bindoff et al., 2019).

Arguably, black carbon (BC) is the most ubiquitous of these allochthonous recalcitrant organic forms. It is produced from the burning of biomass and fossil fuels (Gustafsson et al., 2009). As such the production of black carbon is not usually associated with coastal wetlands or seagrass meadows. This is certainly the case for seagrasses. For mangroves, combustibility has been shown as being unlikely from historical satellite records (Chew and Gallagher, 2018). Although this does not exclude deliberate attempts to ignite a build-up of combustible materials adjacent to the mangroves or dumped inside. For salt marsh, combustion is more common and can be deliberate. However, it appears to be restricted to the tall canopies of relatively dry reed and grass-dominated ecosystems (Nyman and Chabreck, 1995). Nevertheless, given that even the more temperate succulent marshes support a measure reeds and grasses (Prahalad et al., 2020), it may be prudent to test for the autochthonous nature of BC. This can be done either analytically or from historical fire records. In addition to BC, attention has been given to the role of the particulate calcareous carbon fractions (PIC). These have been regarded as the remnants, not of a carbon sink but a carbon source. During their production CO_2_ is formed and expelled into the atmosphere after a chemical redistribution with other dissolved inorganic forms of carbonate and bicarbonate. The extent of which is determined by water body salinity and the atmospheric [CO_2_] (Saderne et al., 2018; Ware et al., 1992).

For allochthonous recalcitrants like BC, the coverage in the literature is still ‘embryonic’. However, PIC measurements are gaining traction where distinctions can be made as a biogenic source in non-geogenic sediments (Saderne et al., 2019). Still, compilations of both BC and PIC remain rare (Gallagher et al., 2020 in press). As a pressing need to increase awareness, we aim to motivate further investigations of blue carbon bias as a stock mitigation service by accounting for the sum of both BC and PIC contributions. To this end, and within the limits of resources, measurements of BC and PIC were made across targeted coastal wetlands. The wetlands were chosen to expect a large bias in traditional stock estimates from the cumulated fractions of BC and PIC to the TOC stocks. To increase generality, the examples were separated by very different geographic and climatic regions, as well as categories of blue carbon ecosystems where the BC content has not been specifically addressed. That is rural temperate salt marshes of Tasmania (Australia), and a tropical urban seagrass meadow within Malaysia. The salt marshes support an abundant calcareous benthic fauna (Prahalad et al., 2020) and fire-resistant succulents, in catchments with a history of fire events. Such fire events are ubiquitous across the large temperate regions of Australia (Attiwill and Adams, 2013). Furthermore, their shallow soils < 1 m are less likely to dilute the BC fraction over older and deeper soils that have seen less extensive fire events (Doerr and Santín, 2016). Stocks are traditionally normalized to 1 m as a reasonable estimate of the depth of oxidation due to the loss of the canopy (Gallagher, 2017; Pendleton et al., 2012). Although it should be noted this BC/TOC soil depth dependence may be confounded by the inability of younger shallower soil profiles to capture the fire event history. The seagrass meadow is the second-largest example across Malaysia that supports abundant calcareous benthic fauna (Shau Hwai et al., 2007). The meadow is affected by both nearby Sumatran peat fires and urban pollution. Furthermore, its patchy seascape and coastal exposure are known to lead to greater amounts of exported allochthonous litter and fine organic clay particles that can elevate BC fractions (Gallagher et al., 2019).

## 2. Study sites

The temperate salt marshes are located in Southeast Tasmania, along the margins of Blackmans Bay and near the small settlement of Murdunna (Fig. 1b). These wetlands were chosen to represent areas known to be affected by two pivotal fire events. The 1967 Black Tuesday bush fires engulfed most of Southeast Tasmania (Tasmania, 1967). A more recent bush fire in 2013 (Hyde, 2013) spread through the catchment and immediate surroundings of Blackman Bay and northern parts of the Forestier Peninsula. The surrounding areas are largely rural with agricultural and forested lands punctuated by small settlements, namely Murdunna, Boomer Bay, Marion Bay, and the village of Dunalley (Fig. 1b). The salt marsh vegetation in the areas is largely comprised of succulent herbs, mainly *Sarcocornia quinqueflora*, mixed with the succulent shrub *Tecticornia arbuscula*, occasionally interspersed with the sedge *Gahnia filum*. The sea rush *Juncus kraussii* is a regular component of the salt marsh vegetation and forms large stands in the case of the Marion Bay salt marsh, the largest marsh in southern Tasmania (Prahalad et al., 2020).

**Fig.1.**
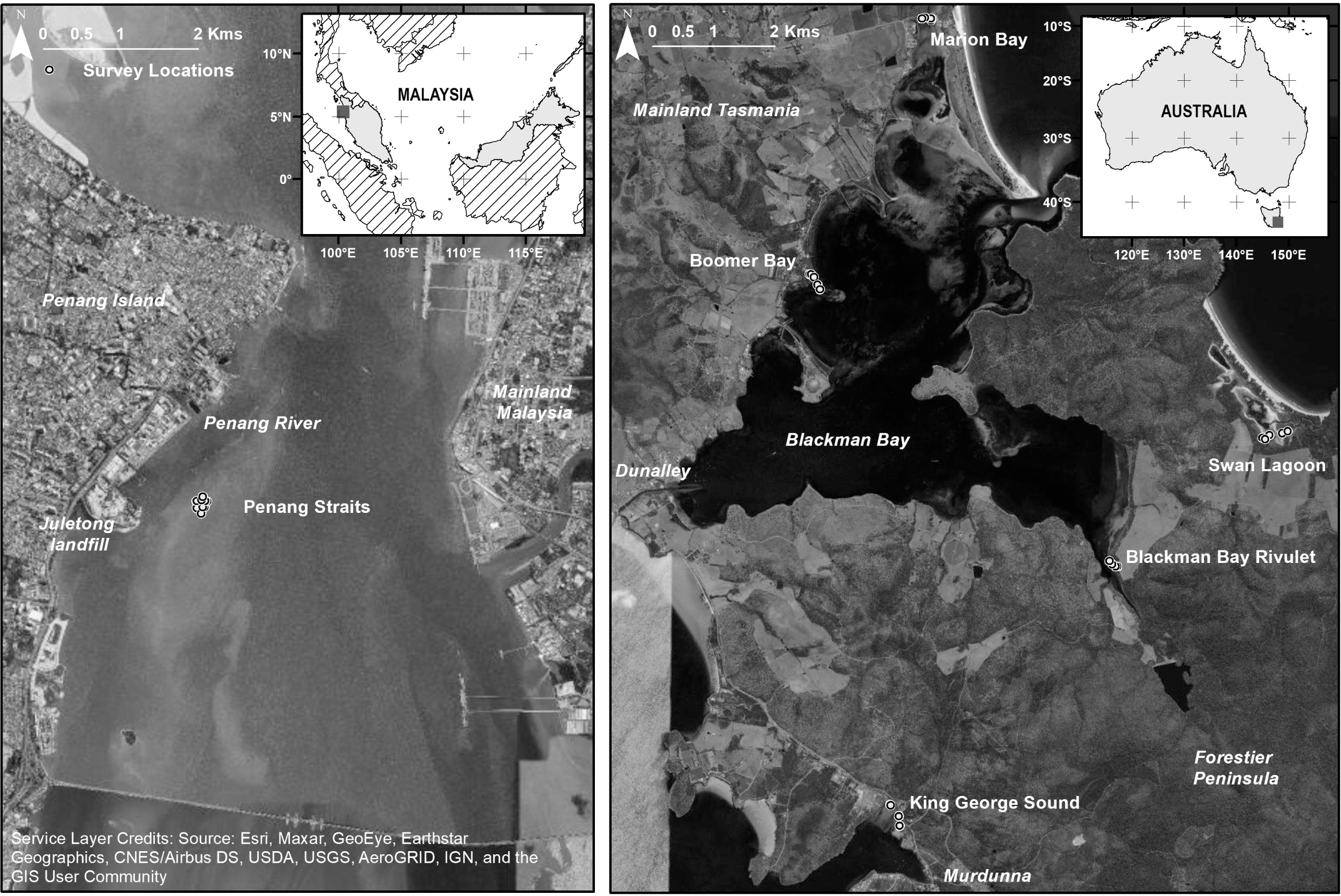
Location of coring sites across the Southeast Tasmanian Salt marshes (Australia) (a), and the Middle bank seagrass meadow within the Penang Straits (Malaysia) (b).

The tropical seagrass meadow occupies Middle Bank within the Penang Straits (Fig. 1a). It represents Malaysia’s second-largest intertidal seagrass meadow (50.7 ha). The meadow is of a patchy configuration dominated by the large leaf *Enhalus acrocoides* mixed with the small-leaved spoon grass *Halodule ovalis*. The Penang Straits harbors a major port, a light industrial complex, and a relatively large urban population density (Chee et al., 2017). Along with industrial and traffic BC combustion products, the region has been regularly affected by the peat fires of Indonesia from the late 20^th^ century (Gaveau et al., 2014). Other possible sources of BC may emanate from the Penang River to the north of the Middle Bank, and the Jelutong municipal waste landfill on the peninsula to the west of the bank (Fig. 1a). Although, supply may be constrained, as the river does not appear to flow directly over the bank (Asadpour et al., 2012). Middle Bank is also largely isolated from the coast with sharp turbidity boundaries of resuspended sediment within the confines of the bank (Lim et al., 2009).

## 3. Methods

### 3.1. Sampling

Salt marsh sampling sites (n = 22) for integrated stock assessments were selected at random towards the seawater edge at low tide at least > 100 m apart within each marsh (Fig. 1b). To identify any confounding of BC dependence with soil stock depths < 1 m, sediment cores were also taken for macro char profiles where the surrounding fires were at their most and least intense across the salt marsh sampling zone. Peaks in char signals were used to date the rates of accretion from the known pivotal fire events (Gallagher and Ross, 2017). Sampling sites for the Middle Bank seagrass meadow were located in the northerly two-thirds of the meadow. Sites were selected randomly at 100 m scales and stratified on the east and west side of the meadow facing the surrounding navigation channels. The final number of sites (n = 15) was determined by the time of exposure at low tide, and an ability to traverse the soft sediments. All sites sampled in both salt marsh and seagrass were assumed to be sufficiently distant as to be depositional independent of each other.

An open-faced ‘Russian Peat Corer’ was used to take 50 cm cores of uncompressed sediment for macro char profiles in vegetated salt marsh soils and stock measurements within seagrass stands. The remaining salt marsh stock samples were exhumed carefully with a PVC pipe and shovel from the base of the soil, where quaternary sands mark the depth of the salt marsh accumulation. The samples where immediately put on ice before transport. The sample cores were mixed well within sealed plastic bags to physically represent the stock variables average; after which a measured volume (≥ 5 cm^3^) was taken for dry bulk density with a cut-off 20 cm^3^ syringe, in the manner of a piston corer. After drying at 60 °C the remaining mixed samples were shaken and sieved through a 200 µm mesh to remove both roots and large shells pieces before further processing for organic matter and elemental carbon analysis (Chew and Gallagher, 2018).

### 3.2. Sample analysis

Subsamples of more than 2 cm^3^ for stock analysis were ground to < 63µm before gravimetric analysis (0.40 g dry weight) for total organic matter (TOM) contents, and calcareous carbon content as PIC (Heiri et al., 2001) using a molecular conversion coefficient (0.273). Particulate inorganic carbon as a mitigation service (PIC_eq_) was adjusted for salinity and atmospheric equilibrium of its released CO_2_ by a coefficient fraction of 0.63 (Ware et al., 1992). All carbon and organic matter contents as dry weight was corrected for salinity (Lavelle et al., 1985). Salinity and pH measurements were taken from the pore water with a handheld refractometer and portable pH meter respectively (Chew and Gallagher, 2018). Black organic matter was isolated after a slow ramping speed to 360 °C according to the protocols devised and tested by Chew and Gallagher (2018). Total organic carbon and BC analysis for Middle Bank sediments were taken from a TOM and BM seagrass sediment calibration curve. The curve was previously constructed from a Malaysian lagoon that included the sediment type and a seagrass genera construct found in Middle Bank (Supplementary material Fig. S1). For the salt marsh samples, TOC and BC were measured directly with a CHN element analyzer after HCl acidification within silver cups at the Central Science Laboratories, University of Tasmania, and after BC was first isolated using the modified of chemothermal oxidation procedure of Chew and Gallagher (2018). In addition to BC contents, their BC stable isotopes signatures were taken from finely ground dry subsamples and analyzed after HCl acidification at the Water Studies Centre, School of Chemistry (Monash University).

The two salt marsh cores taken from the proximity of Marion Bay and King George Sound salt marsh (Fig. 1) were taken for macrochar particle profiles. Before the particles were identified and counted, humic acids and organic matter were removed with KOH followed by H_2_O_2_ on ≈ 2 cm^-3^ of wet sediment and passed through a 100 µm sieved before being washed into a petri dish (Stevenson and Haberle, 2005). The volume was estimated by KOH reagent displacement. The remaining black particles posing as a fractured appearance amongst the now the bleached sediment particles were counted automatically with Image J™ on an enhanced black and white image taken through a magnifying glass (x 5) underlying a white base (Stevenson and Haberle, 2005). A maximum entropy filter was used to isolate the particle images, and any particles in contact with each other were separated with a bone shape function.

### 3.3. Data analysis

Ordinary least squared regressions (OLS) and a one–way analysis of variance for the difference between means (ANOVA), and Student t-test between mean pairs were carried out in Sigmaplot 12™. Where normality was rejected, a Mann–Whitney test replaced ANOVA (Sigmaplot 12™). Comparisons between OLS parameters were determined using an analysis of covariance of their slopes (PAST™). Differences between OLS intercepts of the dependent variable were performed using a Student t-test taken from their standard errors using the 2 degrees of freedom that supports an OLS regression. The precisions as coefficient of variation (CV) were < 10% for all carbon variable: TOC (± 6.3%,n = 14); BC (± 9.0% (n = 4); PIC (± 2.8, n =14). For δ^13^C (± 0.15% (n = 5) the standard error is quoted from other Tasmanian seagrass sediments analyzed at the same laboratory, analyst, and equipment (Gallagher and Ross, 2017). Covariance (n = 5) for char counts was typically 5.3% after repeating the image analysis after remixing the sample identified as the 1968 peak (the numbers ranging between 670 to 750 counts cm^-3^). We recognize that this may be statistically underestimated by not repeating the cleaning stage with a separate subsample. However, the order of magnitude differences in what appears to be the background counts over and the recognizable peaks are not consistent with noise or variance. This contention was supported by the results of a smoothed first derivative with an amplitude threshold of 0.6 and a slope threshold of 0.2 which separated the peaks from baseline variance (O’Haver, 2020) (Supplementary material Peak Detection Template).

## 4. Results

### 4.1. Sediment parameters

All the salt marsh sampling sites had full coverage of succulent plant species (Supplementary material Table S1). Only the coastal wetland strip surrounding King George Sound showed evidence of macrofaunal bioturbation, as observed from the numerous burrows at the surface. Salinities and pH did not significantly correlate with each other (*P* < 0.05, Fig. S2 Supplementary material) and ranged from 1.4‰ to 30.4‰ and 4.83 to 7.83 respectively (Supplementary Material Table S1). Moderate but significant correlations (*P* < 0.05) of pH and salinity with other soil parameters were restricted to peat content with bulk density, soil depth, and other biological soil structures (Supplementary material Fig. S2). All salt marsh data were combined to investigate relationships between variables over landscape scales as one statistical population. Individual carbon concepts across salt marshes displayed equal variances with no differences in their means or median contents (mean TOC *P* = 0.45, mean BC *P* = 0.84, median PIC *P =* 0.67; equal variance TOC *P* = 0.92; BC *P =* 0.60, PIC *P* = 0.15). Middle Bank seagrass sediment pore water salinities were found to be invariant at around 30‰ across the sampling region. Spatial distributions using a kriging model indicated that bulk densities were the smallest and therefore muddier (Tolhurst et al., 2005) towards the NE quarter of the sample area with increasing saltiness (Tolhurst et al., 2005) towards the WSW to SW of the sampling area (Supplementary material Fig. S3). Furthermore, the average BC δ^13^C signatures across all salt marsh sites (−24.86 ± 0.63, CI 95%) was consistent with allochthonous sources. The signatures are typical C3 tree and shrub vegetation signatures (−32‰ to −22‰, average 24‰) and not consistent with the fire susceptible C4 salt marsh reed and cord grasses (−17‰ to −9‰, average −14‰) (Krull et al., 2007).

### 4.2. Organic and inorganic carbon variability

The differences between all salt marsh and seagrass carbon forms were striking (Fig. 2a,b). On average, the salt marsh TOC content was around 35 fold greater than for seagrass. For the remaining BC and PIC_eq_ content variables, this was reduced to around 15 and 4.3 fold respectively from the larger PIC and BC fractions within seagrass sediments. When expressed as a concentration, the differences were reduced considerably. On average salt marsh [TOC] was only 8.3 fold greater than seagrass, with the remaining concepts falling in proportion accordingly (Fig. 2 c,d). This reflects the smaller dry bulk densities found in the salt marsh (0.39 ± 0.10 g cm^-3^, CI 95%) over that of the sandier seagrass sediments (1.50 ± 0.07 g cm^-3^, CI 95%).

**Fig. 2.**
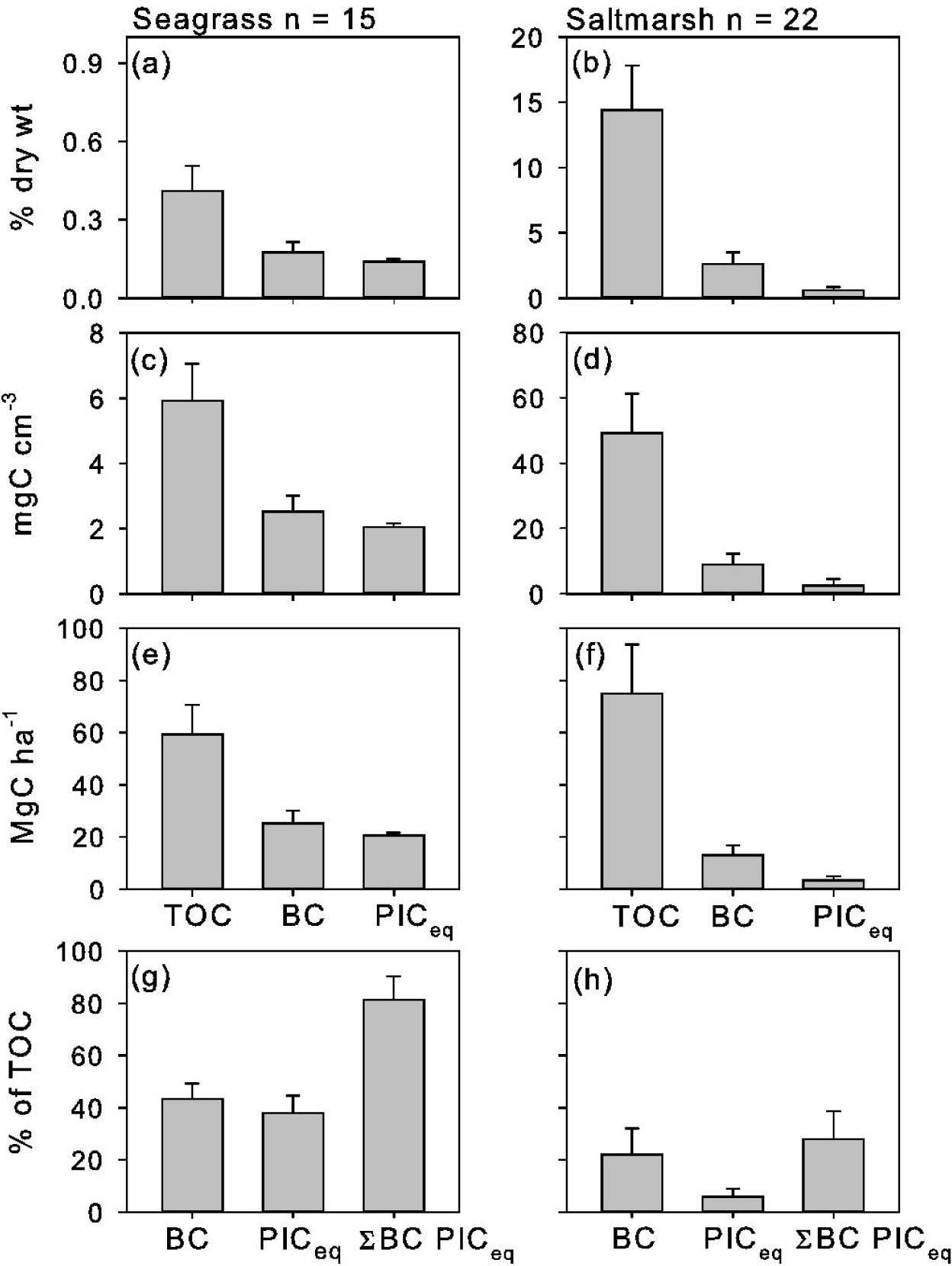
Sediment carbon concepts across the Middle Bank seagrass meadow (left hand charts (a),(c),(e),(g)) and Southeast Tasmania Salt marshes (right hand charts (b),(d),(f),(h)). Error bars refer to confidence limits (CI) 95% about the mean; and TOC, BC, and PIC_eq_ refer to total organic carbon, black carbon, and that fraction of particulate inorganic carbon available for atmospheric exchange in seawater respectively.

In contrast to content and concentration, the TOC stocks between these two ecosystems did not show any significant differences (Fig. 2e,f, P = 0.36 Mann-Whitney Rank Sum Test). The median salt marsh and seagrass TOC stocks were 70.3 MgC ha^-1^ (39.1 to 103.5 as 25% and 75 % quartiles) and 59.5 Mg ha^-1^ (43.2 Mg ha^-1^ to 67.0 Mg ha^-1^ as 25% and 75% quartiles). This convergence was the result of temperate salt marshes’ shallow soils of < 30 cm (Supplementary Material Table S1). Stocks are currently defined by a depth of disturbance at least to 1 m. This is a depth where organic matter is subject to oxidation and remineralization (Pendleton et al., 2012). However, this convergence with TOC stocks between the salt marsh and seagrass did not run to the remaining BC and PIC_eq_ stocks (Fig. 2 e,f). Salt marsh BC stocks diverged to less than half of that found in seagrass (median 13.1 MgC ha^-1^ ± 3.7, CI 95% and 25.2 MgC ha^-1^ ± 4.9, CI 95% respectively, t-test P < 0.001). And salt marsh PIC_eq_ stocks were an order of magnitude smaller than seagrass (a median of 2.3 MgC ha^-1^ 1.2 to 2.4 for respective 25% and 75% quartiles, and a median of 21.0 MgC ha^-1^ 19.2 to 21.3 as 25% and 75% quartiles, respectively, Mann-Whitney Rank Sum Test P < 0.001).

The shallow and variable soil depths across the salt marshes < 1 m implies that their carbon stocks are also a function of depth. However, no dependence was found for salt marsh BC/TOC fractions with soil depth (Fig. 4a), despite a similar average accretion rate over 50 years of deposition of around 0.25 cm yr^-1^ and 0.21 cm yr^-1^. It appeared that any BC/TOC soil depth dependence may have been confounded by an inability for all sites to record all pivotal fires events. King George Sound salt marsh did not record the 2013 fire that destroyed the nearby Dunalley village (Fig. 4b). Overall the average accretion rate across the Southeast Tasmanian salt marsh sites is likely represented as around 0.23 cm yr^-1^. The two accretion sites likely represent the extremes of the sampling region and a type. That is, a site located at narrow open coastal strip over of King George Sound and a site located at extensive Marion Bay saltmarsh within the confines of Blackmans Bay. Given the average [TOC] across all sampling sites of around 49.3 mgC cm^-3^, equates to rates of carbon sequestration across the Southeast Tasmanian sites of around 113.4 gC m^-2^ yr^-1^. In terms of carbon sequestration services corrected for BC deposition, and BC and PIC_eq_ equate on average to around 88.5 gC m^-2^ yr^-1^ and 81.7 gC m^-2^ yr^-1^ respectively, a total mitigation bias of around 18.3%.

In summary, irrespective of the similarity between the tropical seagrass meadow and temperate salt marsh TOC stocks (Fig. 2g,h), mitigation corrections for BC and PIC were very different. For seagrass, the BC/TOC fraction was around 43.3% ± 5.9% (CI 95%) with nearly matching constraints from PIC_eq_ of 38% ± 6.6 % (CI 95%). In total, there was 81.3% ± 9.0 (CI 95%) overestimate in sedimentary carbon stocks. This is in contrast to the salt marshes, where the smaller but moderate BC/TOC fraction of 22.0% ± 10.2% (CI 95%) was supplemented by a significant but minor PIC_eq_/TOC fraction of 6.0% ± 3.1 (CI 95%). Nevertheless, taken together this is a significant overestimate of more than a third of carbon stocks as a mitigation service (28.0% ± 10.8% CI 95%). For the average salt marsh carbon sequestration services, which were independent of soil depth the bias was less extensive (18.3 %).

### 4.3. Dependence of carbon concepts with total organic carbon

Ordinary least squares regressions across the seagrass meadow produced a strong positive correlation of sedimentary BC contents with its TOC counterpart closely trending from the origin (Fig. 3a r = 0.84, □ = 0.99). In contrast, the salt marsh BC–TOC content regression model displayed only a moderate correlation, with good power but with poor predictive value (Fig. 3c r = 0.62, □ = 0.88). The poor prediction is reflected in the relatively larger variance around the regression line. However, this result may have been leveraged by a single large sample value (Fig. 3c ■, Cook’s distance = 1.2). After the removal of this likely outlier, the resultant regression became weaker and exhibited low power (r = 0.39, □ = 0.43). For the seagrass meadow’s sedimentary PIC contents, significant dependence on its TOC counterpart was found (*r* = 0.83, □ = 0.99), but with relatively little variation in PIC with TOC contents (Fig. 3b). This invariance with TOC content was similar across the salt marsh (Fig. 3d).

**Fig. 3.**
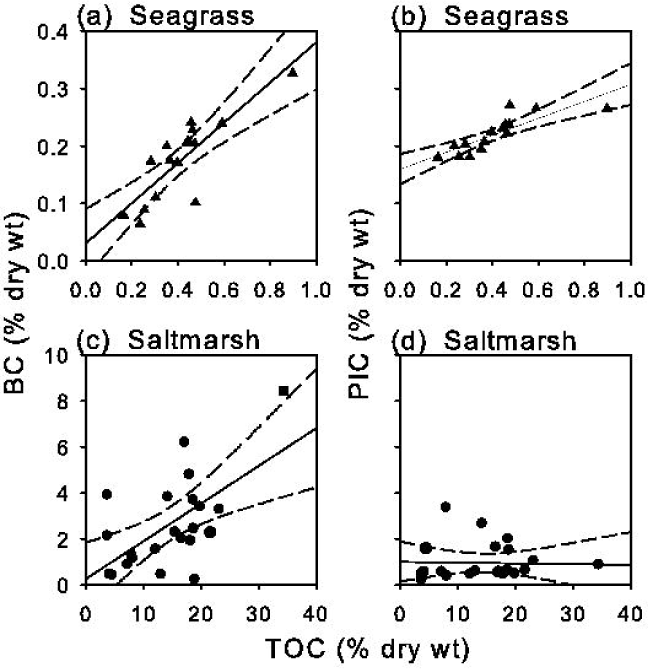
Above (a), the relationship between the fraction of black carbon (BC) within the total organic carbon (TOC) pool with soil depth across Southeast Tasmanian salt marshes, the dotted lines refer to 95% confidence limit boundaries either side of an ordinary least squares regression. Below (b) illustrate the macro char profiles down the soil profiles at the northern and southern boundaries of the sampled salt marshes. 1968 and 2013 mark the positions of the two pivotal fires of the period.

## 5. Discussion

Measuring and comparing values of carbon concepts (e.g., stocks, concentrations, and contents of TOC, BC, and PIC) across underrepresented blue carbon ecosystems and environs are useful measures of variance towards global assessments. However, additional insight and predictive power are also possible by their relationship with each other (Chew and Gallagher, 2018; Gallagher et al., 2019). The following discussion examines the statistical ranges and relationships between those carbon concepts, as a measure of their importance and insights on the nature of the supply of BC and PIC variance

### 5.1. Black carbon sources

It has been argued that the structure of BC–TOC regressions is the result of the addition of two possible supply paths, namely, aeolian deposition and soil washout. With aeolian deposition, sedimentary contents of BC across a meadow or wetland is independent on its TOC. The extent of residual variance in BC is likely affected by different accretion processes that may vary over larger distances than those areas restricted across embayments (Chew and Gallagher, 2018; Gallagher et al., 2019; Sun et al., 2008). Whereas sedimentary BC supplied from soil washout increases with TOC, where any the BC intercept represents the underlying remnant of BC invariance from additional aeolian deposition (Chew and Gallagher, 2018). For the Tasmanian salt marshes, the relative invariance of BC with TOC appears to be consistent that a supply of BC dominated by aeolian deposition (Fig. 3c). As BC emitted to the atmosphere is undiluted by soil organics aeolian deposition would be expected to elevate the BC/TOC sedimentary fractions (22% ± 10.2%, CI 95%, and median 14.7%) over a salt marsh supplied largely from BC diluted with soil organics.

Comparisons with other salt marshes do not appear to be available. Although there is data available for an autochthonous refractory carbon (C_rf_) for two humid subtropical climate salt marshes across the USA and two temperate oceanic climate salt marshes across the Iberian Peninsula (Leorri et al., 2018). The authors consider that this fraction contained a pyrogenic had been washed in from the surrounding landscape along with other geological refractory sources. Although, it appears that the analytical method was originally used for isolating BC. Nevertheless, they reported C_rf_ /TOC fractions values were still smaller than our temperate BC fractions, ranging from 6.4% to 13.1% and 13.1% to 17% as dry wt for the USA and Iberian pairs, respectively. All things being equal, comparisons between the above salt marsh examples alone, would suggest that aeolian transport from surrounding fires may the vector that is largely responsible for the elevation of BC/TOC fractions. It was noted that the bush fires of 2013 immediately surrounding the Tasmanian salt marsh were quickly deposited as large macrochar particles. This was not only felt by the author (JBG) at the time of the fire, and also observed as heavy deposits washed up on the beaches and the local seagrass surface sediments after the 2013 fire (Supplementary material Fig. S4). It is likely the dilution of sedimentary BC contents was great within the Tasmanian salt marsh. The average Tasmanian salt marsh carbon sequestration rate (113.4 gC m^-2^ yr^-1^) is close to half of the global average (244.7 g Cm^−2^ yr^−1^) (Ouyang and Lee, 2013). Interestingly, rates of TOC sequestration across the Tasmanian salt marshes were notably larger than the tidal marshes of the neighboring Australian mainland, across the state of Victoria (86.86 gC m^-2^ yr^-1^ ± 4.79 SE) (Macreadie et al., 2017). The differences appear to be largely the result of a higher TOC content across the Tasmanian salt marsh (14.41 % dry wt ± 1.63 SE and 7.86 % dry wt ± 0.59 SE respectively). It is unlikely that the larger content comes from BC additions, as Victoria has not been immune from pivotal fire events (Forest Fire Management Victoria, 2020).

In contrast with the Tasmanian salt marsh, the BC contents across Middle Banks’ tropical seagrass appear to show an unexpected dependence with TOC (Fig. 4a). This is consistent with a supply of BC largely from soil or adjacent sedimentary washout (Chew and Gallagher, 2018; Gallagher et al., 2019). It appears that Middle Bank may not have been as isolated from the shoreline as first hypothesised. Although the small BC positive intercept suggests an additional aeolian deposition that elevated the meadows’ BC/TOC fraction. Interestingly, the proportional response of BC with TOC is statistically identical to that found across a rural meadow dominated by the same seagrass species (Enhalus spp.) at Limau Limauan (*P* = 0.64, reanalysed from supplementary material attached to Gallagher et al., 2019). Although overall, Middle Bank supported a larger BC/TOC fraction than Limau Limauan (26.1% ± 4.9 CI 95%). This may be a combination of a smaller fraction of aeolian deposition based on a statistically smaller BC intercept (*P* = 0.003 of the two intercepts being equal) and the extent of black carbon pollution between these regions. Limau Limauan is adjacent to the shoreline and located within a North Borneo marine park, a region located within the penumbra of the Southeast Asian BC hotspot (Chew and Gallagher, 2018; Permadi et al., 2018). Nevertheless, the differences between Limu Limaun and the Middle Bank seagrass meadows did not reflect the larger categorical differences in BC emissions between their respective regions’ (Permadi et al., 2018). The apparent normalization of this difference may reflect their different tidal niches. Middle Bank is intertidal and Limau Limaun is subtidal. It has been noted that tidal exchange can remove BC from intertidal wetlands to coastal waters as it dissolved after conditioning by a slow microbial attack (Dittmar et al., 2012). Indeed, such a process is also consistent with the smaller concentration of BC from what appears to be the more extensive 1967 ‘Black Tuesday fire’ over the surface peak from the 2013 fire (Fig. 3a).

**Fig. 4.**
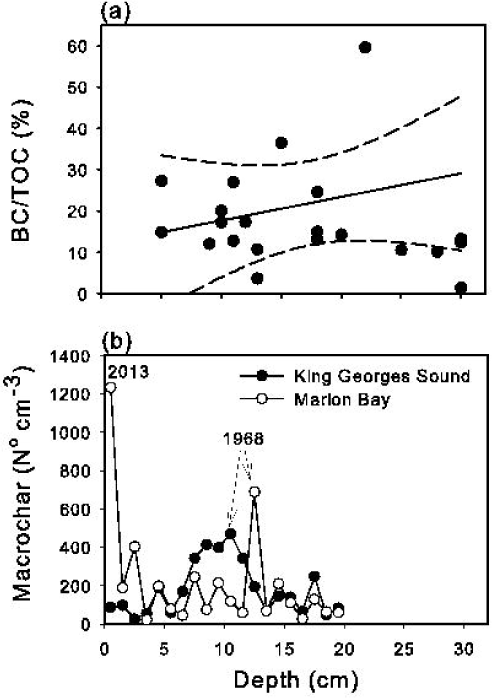
The relationship between sedimentary total organic carbon (TOC) with black carbon (BC) and particulate inorganic carbon (PIC). Charts across the top (a) and (b) represent Penang’s Middle Bank’s seagrass meadow, and across the bottom (c) and (d) represent Southeast Tasmanian salt marshes. The dotted lines refer to 95% confidence limit boundaries either side of an ordinary least squares regression, and the point marked as a ■ in Figure 3c is an outlier that possesses large leverage on the regression’s parameters (see text).

### 5.2. Particulate inorganic carbon variance

The relative invariance of PIC with TOC content, more so for salt marsh than the seagrass meadow, may simply reflect parameters determined by their different canopy as a calcareous faunal niche. Indeed, extensive and relatively constant fractions of PIC with near-identical TOC stocks have also been reported for Merambong, another Malaysian intertidal coastal seagrass meadow that supports an *Enhalus spp*. (Rozaimi Jamaludin et al., 2017). However, PIC stocks were more than double that of the Middle Bank meadow. The reasons for this difference are unclear, other than due to the difference in sample preparation. There was no indication that there has been a need to separate any larger calcareous pieces < 2 mm that would otherwise elevate the PIC content (Rozaimi Jamaludin et al., 2017).

### 5.3. Carbon stocks and sequestration as a mitigation service

The results indicate that carbon stock services in mitigating greenhouse gas emissions may not be significant for tropical intertidal coastal seagrass meadows, which appear to support an abundant calcareous and occupy BC hotspots. In the same vein, current carbon stock mitigation for temperate salt marshes estimates may be required to be reduced by up to third (28.0% ± 10.8% CI 95%). This is largely by their BC contributions in relative terms and absolute terms by their shallow soil depths < 1 m. How these fractions are used to correct for carbon sequestration mitigation bias will depend on the extent of organic mineralization over the long term. Traditionally, this has been assumed to have largely ceased near the surface after 1 or 2 years of deposition (Cebrian, 2002). However, this assumption has recently been challenged There is evidence to suggest that mineralization within mangroves sediments continues after 100 years and that 50% of the TOC, excluding BC, may be lost from seagrass sediment horizons within a 100 years of deposition (Chuan et al., 2020; Maher et al., 2017).

### 5.4. Other limitations

It has been accepted that despite dissolution (Dittmar and Koch, 2006) BC remains largely unmineralized over climatic scales (Kuzyakov et al., 2014). In contrast, accounting carbonaceous mitigation services are not as straight forward. Current blue carbon conceptual models caution the presence of geogenic carbonates from the catchment (Saderne et al., 2018) or indeed the remains of coral sands. Their production and associated emissions of CO_2_ have been produced outside the ecosystem. Such a geogenic constraint is not necessary for the Middle Bank and the Tasmanian salt marsh sediments. It would then seem that only the ecosystems’ biogenic carbonates play a role as an ecosystem carbon source. However, a more considered ecosystem service logic could argue for a different set of considerations. These are the services that account for the source and stability of all carbonates relative to those services, and emergent services within a replacement non-vegetated ecosystem. For example, the dissolution of coral rubbles by anthropogenic ocean acidification outside vegetated canopies (Eyre et al., 2014; Sulpis et al., 2018). Such a process adds to the coastal dissolved inorganic carbon sink (DIC) pool which turns over at millennial scales (Maher et al., 2018). The removal of this carbon sink service could be conceivable be constrained by the presence of a vegetated coastal canopy. Canopy photosynthesis is known to elevate its water column pH (Koweek et al., 2018; Krause-Jensen et al., 2016) and would thus constitute a reduction in mitigation services over the non-vegetated alternative. Additional consideration may also be required on the origin of the ecosystem’s biogenic carbonates. The production of calcium carbonates as part of a photosynthetic carbon concentrating mechanism has been noted for some seagrass (Enriquez and Schubert, 2014) and seaweeds (Borowitzka and Larkum, 1987). As carbonate precipitate the emission of CO_2_ within the lacunae their thalli and leaves, are in large part, recycled to RuBisCO for carbon fixation. Consequently, their presence is the remnant of an additional organic carbon sink and not source. Indeed such plant carbonates can be a major contributor to carbonaceous sediments (Enriquez and Schubert, 2014; Perry et al., 2019). For salt marsh plants, this process as far as we are aware has not been tested.

Other mitigation constraints previously discussed in the literature concern the dissolution of biogenic carbonates after the loss of the canopy. Oxidation and dissolution of carbonates lead to alkalization and may reduce the water columns’ *p*CO_2_ from sediment organic oxidation (Howard et al., 2018). On the other hand, there are whiting events associate with tropical coastal waters (Broecker et al., 2000). These calcium carbonate precipitates have been seeded by sediment resuspension. However, whiting events after the loss of seagrass canopies have not yet been reported. Nevertheless, such events could conceivably increase the water columns *p*CO_2_ during wind resuspension from the loss of a seagrass canopy (Gallagher et al., 2020 in press).

## 6. Conclusions

It was found that allochthonous BC was likely to be a major contributor to the TOC sedimentary stocks (around 43.3%) for coastal seagrass meadows that occupy BC hotspots, such as Middle Bank (Peninsula Malaysia). Although, the extent of this fraction may be reduced by circumstances of intertidal flushing of its dissolved BC. Furthermore, only a minor fraction of the carbon stock remained for Middle Banks’ ability to mitigate greenhouse gas emissions when the BC constraint was pooled a similar fraction of the biogenic PIC_eq_ (around 38%). In contrast, the temperate salt marshes subject to pivotal fire events likely supported a moderate allochthonous BC fraction of around 22% of its TOC stocks, with minor contributions from the PIC_eq_ (around 3.2%), such as Southeast Tasmania. This BC fraction appears to be elevated over another less temperate salt marsh with a similar allochthonous carbon concept. The reasons are unclear, other than salt marshes in Tasmania supports a shallower sediment profile, lower than the average global rate of carbon sequestration, and has been impacted by a nearby large forest fire events. These appear to have supplied BC largely undiluted by aeolian deposition. These examples of extremes if not ubiquitous, by themselves represent areas that are global extensive. Either way, it would be prudent to address the sources, recalcitrance, and function of particulate carbon types to address what can be considered bias in carbon stocks, and sequestration mitigation assessments. This will assist in avoiding both local and global overestimates and ensure that greenhouse gas emitters not to exceed their capacity within a carbon sink management scheme.

We also add a word of caution. The arguments for subtracting biogenic PIC from the blue carbon mitigation equation is not as unequivocal as BC. The risk could divert carbon credit resources from restoration or preservation for ecosystems that support biogenic PIC fractions equal to or greater than organic carbon stocks. This would also omit their less tangibly valued but real additional ecosystem services, such as biodiversity, fish nurseries, coastal protection, and so on (Barbier et al., 2008). More research is needed away from the traditional mass balance of inorganic and organic elements to one under the umbrella of mitigation carbon mitigation services and relative to the canopy and non-vegetated alternatives.

## Supporting information

Peak detection template

## CRediT authorship contribution statement

**John Barry Gallagher**: Conceptualization, Validation, Formal analysis, Investigation, Writing - original draft, Writing - review & editing, Visualization, Resources. **Vishnu Prahalad**: Visualisation, Resources, Writing - original draft, Writing - review & editing, Funding acquisition. **John Aalders**: Investigation, Resources, Writing - review & editing.

## Acknowledgments

We acknowledge the support of Jeff D Ross of the Institute of Marine and Antarctic Studies, University of Tasmania for facilitating the authors’ position of Adjunct Researcher, and the support of the Centre of Marine and Coastal Sciences, Universiti Sains Malaysia.

