## Supplementary Materials for "Inorganic and black carbon hotspots constrain blue carbon mitigation services across tropical seagrass and temperate tidal marshes"

**Introduction**

This document contains additional information cited to support the main articles' contentions, and uncited additional matrix and species parameters, for both context and future regional and global comparisons. These include: 1) additional figures showing the quality and relationship of calibrations used in the study; 2) data used in the construction of figures used in the main articles along with their site coordinates are tabulated; 3) additional ecological parameters within sampling sites; 4) relationships cited within the main article between available ecological parameters required to focus on the important variables that could otherwise constrain the interpretation of the article.

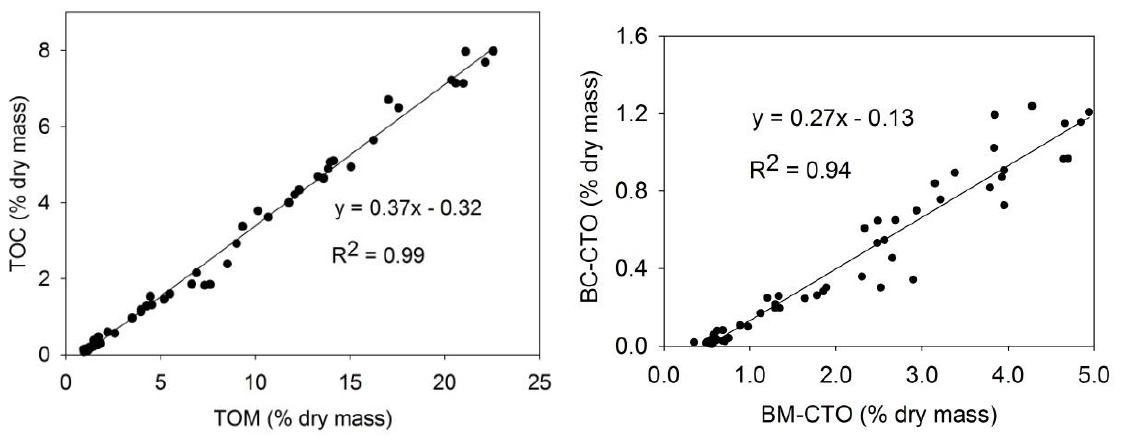

Figure S1. Total organic carbon (TOC) from total organic matter ((TOM) as loss of ignition, black carbon (BC) after chemo thermal combustion separation (CTO) (n =55) across the seagrass meadows of the lagoon Salut–Mengkabong (Sabah Malaysia) (Chew and Gallagher, 2018). The figures cited by (Chew and Gallagher, 2018), and taken directly from Chew (2018) MSc thesis <http://eprints.ums.edu.my/id/eprint/25165> with permission from Chew.

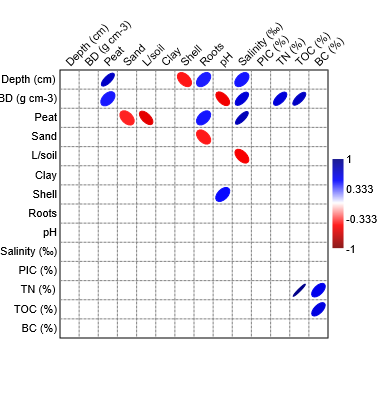

Figure S2 illustrates an ellipse linear Pearson plot of significant correlations r (*P* < 0.05) between the Southeast Tasmanian salt marsh sedimentary parameters and carbon variables (the matrix was constructed in PAST™). BD is dry bulk density, L/soil s the loam fraction, PIC- particulate inorganic carbon content, TN is the total nitrogen content (uncorrected for any ammonia sorption) TOC is total organic carbon content, and BC is black carbon content after isolation by chemothermal oxidation (Chew and Gallagher, 2018).

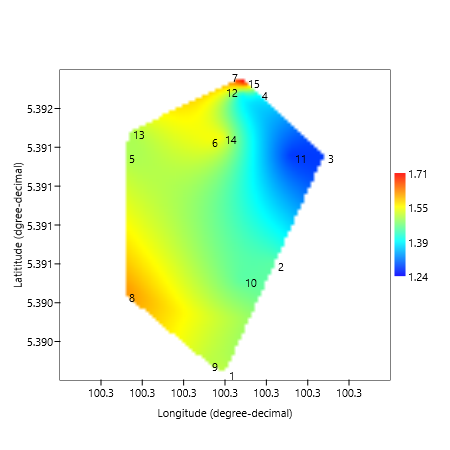

Figure S3 a kriging contour of sediment dry bulk densities (g cm^-3^) at sampling stations across the Middle Bank seagrass meadow, Penang straights, Malaysia (constructed in PAST™ ). Note the generally higher bulk densities on the west side of the bank facing the navigation channel between Penang Island and Middle Bank (Fig.1). Lower dry bulk densities in less consolidated muddier sediments noted during traversing the bank, occupied the eastern side of the middle bank facing the main body of the Penang straights and the Malaysian Peninsula (Fig. 1).

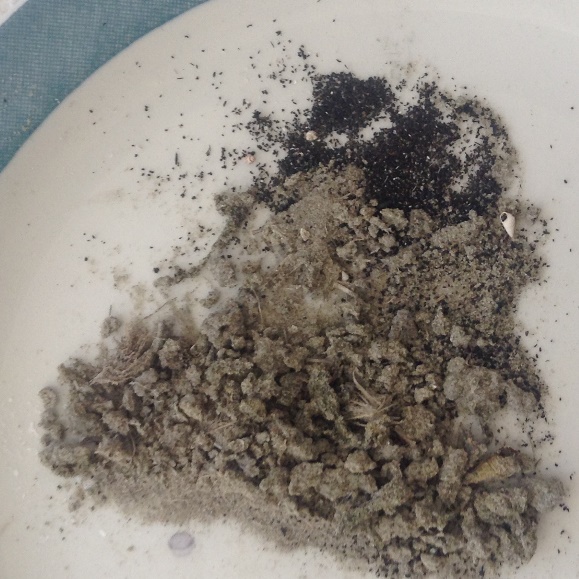

a

b

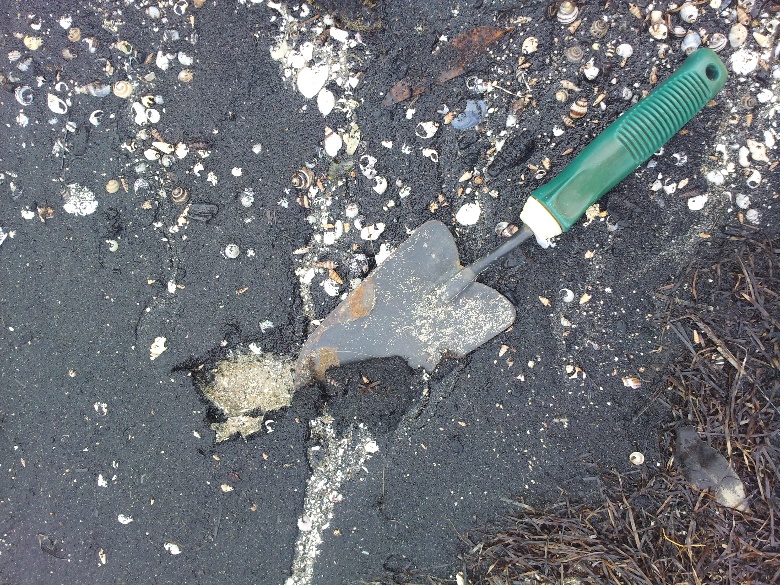
Figure S4 a month after the 2013 surrounding bushfires of the remnants of black carbon deposited within the surface sediments of the bay’s intertidal seagrass meadows (a, -42.904627° S and 147.819515° E) and washed onto the beaches surrounding Blackmans Bay (b, -42.840085° S and 147.853381° E).

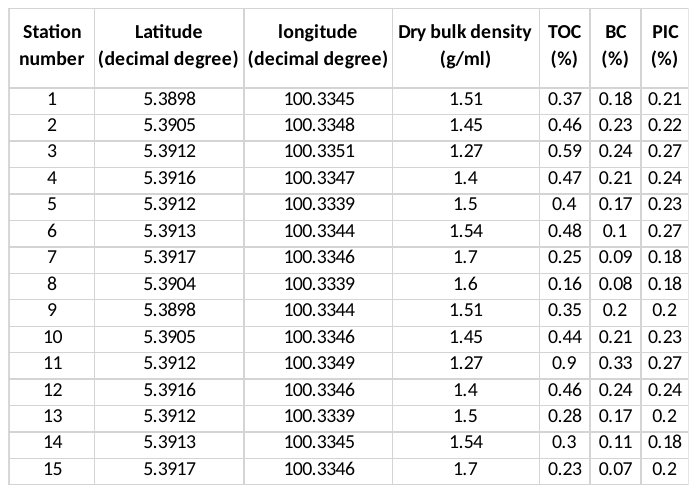
Table S1 the Middle bank seagrass (Penang, Malaysia) sediment variables and locations. Note that pore water salinities were relatively constant across the meadow (30‰) as measured with a portable refractometer at stations 1, 2, 6, 8, and 15. All carbon contents we corrected for salt from their water content, obtained during dry bulk density measurements, and salinity of 30‰.

Table S2 Southeast Tasmanian (Australia) salt marsh sediment and plant variables, with sampling site locations

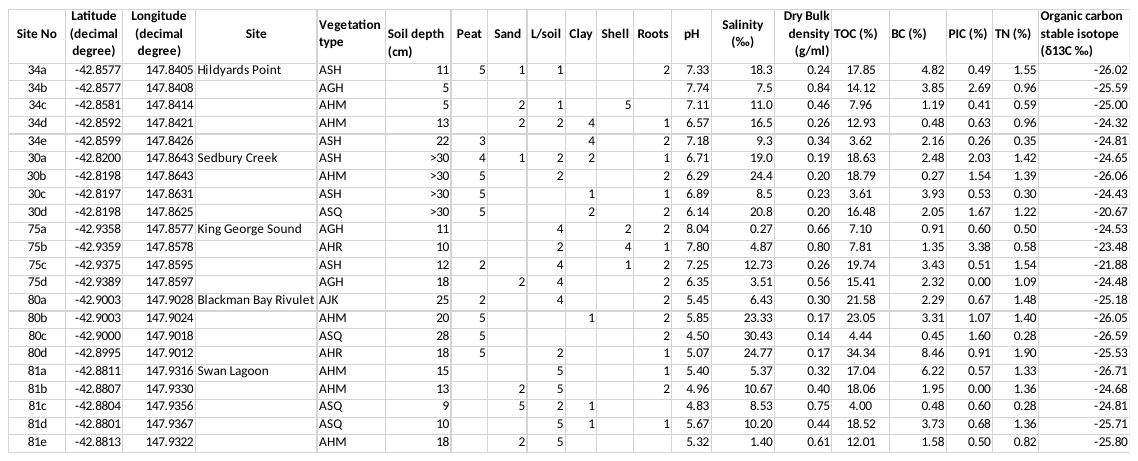

Vegetation codes: AGH = *Graminoids* and herbs; AHM = mixed herbs; AHR = herbs and rushes; AJK = *Juncus* *kraussii*; ARH = rushes and herbs; ASH = shrubs and herbs; ASQ = *Sarcocorni*a *quinqueflora*; ASS = *Sarcocornia and Samolus*.

Composition as peat, sand L/soil (loam), clay, shells, roots for the surface 10cm: 0 = <1%; 1 = 1 - 10%; 2 = 11 - 25%; 3 = 26 - 50%; 4 = 51 - 75%; 5 = >75%
